# Hidden in Plain Sight: A Novel Symbiotic Haplosclerid Sponge Species Revealed by its Mitochondrial Genome

**DOI:** 10.1101/2025.07.08.663769

**Authors:** Momin Ahmed, Jan Vicente, Russell T. Hill, Dennis V. Lavrov

## Abstract

An unusual third mitochondrial genome was discovered in the genomic data from the symbiotic association between two sponges, *Haliclona plakophila* (Haplosclerida, Demospongiae) and *Plakortis symbiotica* (Homoscleromorpha). This mitogenome exhibited an unusually high GC content, lacked the *cox2* gene, and could not be reliably positioned among animals based on phylogenetic analysis of the sequence data. Nevertheless, conserved gene clusters and the presence of *atp9* indicated that this species is a sponge, while phylogenetic analysis using *18S* and *28S* places it in the order Haplosclerida of Demospongiae. Since no tissue of the cryptic sponge was found in the investigated association, its relationship with other sponges remained unclear. Therefore, we sampled 10 additional samples assumed to be *H. plakophila – P. symbiotica* associations in the field to search for the mysterious third sponge. PCR analysis revealed that one of the collected samples contained the novel sponge, but not *H. plakophila*. A follow-up morphological investigation revealed that this sponge, **Haplosclerida sp. nov**., while superficially indistinguishable from *H. plakophila*, differs in skeletal organization and spicule composition. Here, we report the unusual mitochondrial genome of the new species, along with a phylogenetic analysis using *18S* and *28S* sequences, and demonstrate that **Haplosclerida sp. nov**. belongs to a new clade of Haplosclerida. We also report the mitogenome of *Xestospongia deweerdtae*, another haplosclerid species described in association with multiple *Plakortis* species, including *P. symbiotica*. Our results support multiple origins of Haplosclerida-Plakortis symbiosis and open new avenues for its investigation.

## Introduction

Sponges (phylum Porifera) are sessile, benthic organisms that often compete with other organisms, including other sponges, for seabed real estate to grow on. Consequently, many sponges have evolved strategies to cope with such pressure; symbiosis is one among them. Symbiotic associations between sponges have been reported as far back as the Challenger expedition to the Indian Ocean, where Sollas (1888) observed one sponge overgrowing and penetrating the tissue of another. Other observations in crowded cave environments showed species growing on the same substrate, intimately in contact, or one sponge epizootically growing atop another, sometimes completely engulfing it without suffocating it (Rützler, 1970; Sarà, 1970). While the nature of these associations is not well understood, several potential benefits to the sponge partners have been proposed (discussed in Wulff 2008, Wulff 2011, Wulff 2021). One of them, the ability to share chemical defenses against predation, was experimentally demonstrated by Ramsby *et al*. (2012). This study showed that the chemically defended *Amphimedon erina* overgrew *Geodia vosmaeri* (a palatable species to predators), protecting it from the spongivorous sea star *Echinaster sentus*. When sharing such defenses, sponges may be able to expand into environments where they typically would be removed through predation (Wulff, 2008).

Recently, Vicente *et al*. discovered several novel two-sponge associations in the Caribbean (Vicente *et al*., 2014; Vicente, Zea, and Hill, 2016) between two *Plakortis* species: *Plakortis symbiotica* and *Plakortis deweerdtaephila*, and two representatives of the demosponge order Haplosclerida: *Xestospongia deweerdtae* and *Haliclona (Halichoclona) plakophila* (Figure 1A). *P. symbiotica* was found to be associated with both haplosclerid species, while *P. deweerdtaephila* seems to associate exclusively with *X. deweerdtae*. Except for *X. deweerdtae*, none of the sponges were found in freeliving condition outside of symbiosis. Symbiotic *X. deweerdtae* has a significant decrease in the size of their strongyles (22% width, 30% length) compared to the free-living form, which suggests they invest less energy in skeleton synthesis when in association.

**Figure 1.**
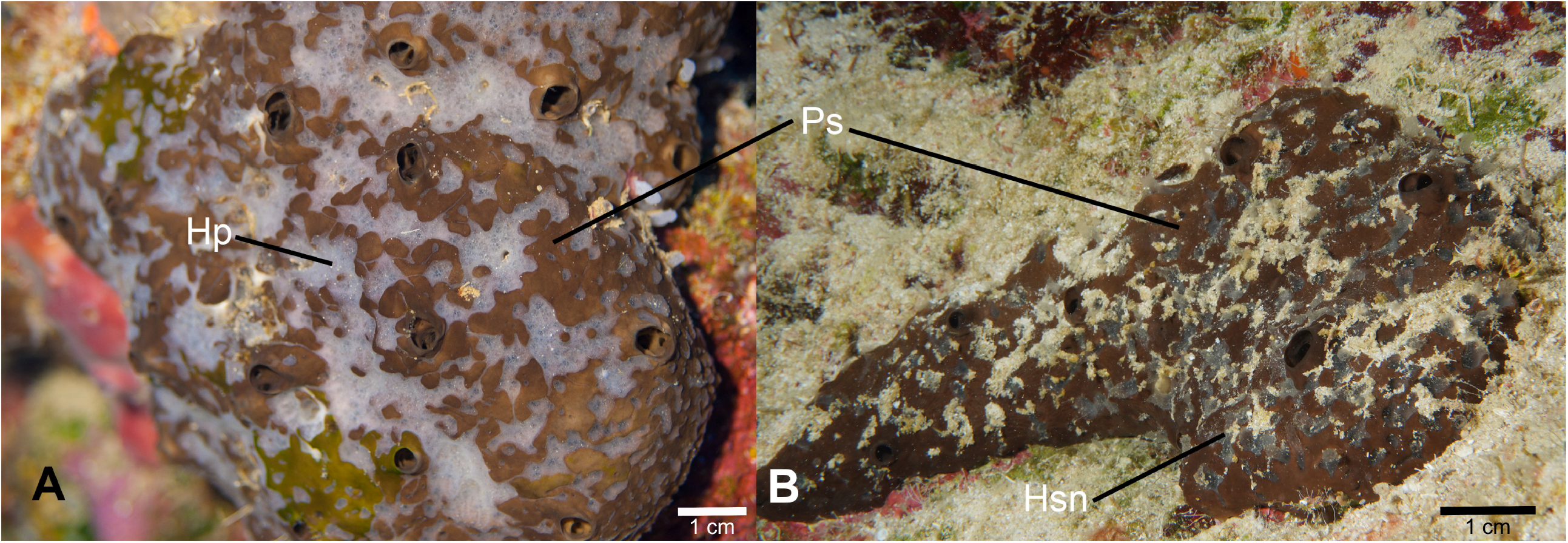
The two symbiotic associations involving *Plakortis symbiotica*. A) Holotype USNM1254650 specimen of *P. symbiotica* basibiont (Ps, brown) with the associated *Haliclona plakophila* epibiont (Hp, bluish-white). B) Specimen BPBM-C1914, Ps with associated **Haplosclerida sp. nov**. (Hsn, bluish-white).

Our recent study (Lavrov, Turner, and Vicente, 2025) used low-coverage sequencing of *Haliclona plakophila* that had been dissociated from their symbiotic partner to obtain a complete mitochondrial genome of that species. Surprisingly, the resulting assembly contained three mitochondrial genomes (mitogenomes). Two of these were expected: one was determined to belong to clade B of Haplosclerida and was subsequently described in the same publication as belonging to *Haliclona plakophila*. The second was reconstructed as an 18.3-kbp-long circular molecule with the typical gene content and order reported in *Plakortis* species (Wang and Lavrov, 2008) and shared 100% identity with previously reported *Plakortis symbiotica* partial sequences. The third mitogenome, however, appeared to be quite unusual. Reconstructed as a 17.2-kbp-long circular molecule, it had a highly GC-biased nucleotide composition and lacked *cox2*. The unusual composition of the novel mitogenome initially made it unclear what type of organism it belonged to; phylogenetic analysis based on both mitochondrial nucleotide sequences and inferred mitochondrial protein sequences proved uninformative, producing a long and unstable branch in phylogenetic trees. However, the presence of the conserved gene block *atp8*->*atp6*->*cox3* (compared to ancestral bilaterian gene order, Lavrov *et al*., 2005) indicated that this mitogenome is of animal origin, while the presence of *atp9* narrowed it down to the phylum Porifera (Lavrov, 2007).

We initially hypothesized that this mitogenome may be a third member of this symbiotic association, possibly as a parasite (Ahmed, 2024). We also speculated that this third mitogenome might belong to *X. deweerdtae*, as this species is also known to form symbiotic relationships with the same basibiont. To resolve this cryptic mitogenome’s identity and phylogenetic position, we collected ten additional samples of the *H. plakophila — P. symbiotica* association and used PCR amplification to identify the novel mitogenome within the associations. We also sequenced total DNA from a previously collected sample of *X. deweerdtae* and described its mitogenomes as well as nuclear rRNA.

While all the samples of *H. plakophila — P. symbiotica* association appear indistinguishable in the field, PCR amplification using species-specific primers revealed that one of them (BPBM C1914) contained the unknown third sponge. Surprisingly, the PCR analysis did not find the presence of *H. plakophila* in this sample. The sequence of the unknown sponge was different from that of *X. deweerdtae*. Hence, we propose that this association represents a novel symbiosis between *P. symbiotica* and an as-yet unknown haplosclerid sponge. Here we present a brief morphological description of the new sponge, which we call **Haplosclerida sp. nov**., along with the description and comparison of its mitogenome, and an analysis of its phylogenetic position. We also show that symbiotic relationships between *P. symbiotica* and Haplosclerida evolved independently at least three times.

## Methods

### Sample Collection and Preservation

Samples of the *Haliclona plakophila – Plakortis symbiotica* associations were collected by SCUBA at a depth of 25-33 meters in the La Parguera region, Puerto Rico, in July of 2024 at three sites (Black wall (N 17 ° 53.091’, W 67° 0.921’), Hole in the Wall (N 17 ° 53.076’ W 67° 1.332’) and Fallen Rock (N 17 ° 53.910’ W 66° 56.772’). A total of 10 samples were collected and stored in 95% ethanol and RNA later. Specimens were collected under scientific collection permit number 2024-IC-046 issued by the Department of Natural Resources (covering the period of July 12, 2024, through July 31st, 2025) and deposited at the Bernice Pauahi Bishop Museum in O?ahu, USA. Collection information and permits for samples of *X. deweerdtae* are described in Vicente *et al*A., 2014, Vicente *et al*A., 2016.

### Spicule Preparation and Sectioning of the Ectosome

Sponge pieces (1 cm^3^) containing both ectosomal and choanosomal tissue were removed from 95% ethanol and digested in household bleach (5 – 6 % sodium hypochlorite solution) for 30 minutes at 50°C. Spicules were left to settle, and bleach was discarded. Spicules were then rinsed two times with distilled water to remove any remnants of the bleach solution, and the water was changed to 95% ethanol for long-term storage. A few drops of the spicule suspension were added to a stub, air-dried, coated in gold for 45 seconds, and imaged under a Hitachi S-4800 FESEM Scanning Electron Microscope (SEM) at the Biological Electron Microscope Facility at the University of Hawai’i at Mānoa. Spicules were also observed under light microscopy, photographed, and measured using ImageJ (Abràmoff, Magalhães, and Ram, 2005). The lengths and widths (L x W) from thirty oxeas were measured; for sigmas, length, width, and thickness (L x W x T) of 10 to 15 spicules were measured following Figure 1 in Van Soest (2017). Thick sections of the ectosome were hand-made directly on the voucher specimen fixed in 95% ethanol by carefully lifting the **Haplosclerida sp. nov**. epibiont pieces off the *Plakortis symbiotica* tissue with a scalpel. The sponge pieces were then placed on a microscope slide, supplemented with additional ethanol, and mounted with a cover slip. Ectosomal sections were examined and photographed under light microscopy. After processing, sections were returned to the ethanol jar with the voucher specimen.

### Genome Sequencing, Assembly, and Annotation

Total DNA of **Haplosclerida sp. nov**. and *Xestospongia deweerdtae* was extracted using a phenol-chloroform method modified from Saghai-Maroof *et al*. (1984). Genomic DNA libraries were prepared for **Haplosclerida sp. nov**. using the NEBNext Ultra II Library Prep Kit according to the manufacturer’s protocol, and for *X. deweerdtae* genomic DNA/cDNA/BAC PCR-free library was constructed at the DNA Facility, Office of Biotechnology, Iowa State University. Whole-genome sequencing was performed at the DNA facility, using the Illumina NovaSeq 6000 platform with 150-cycle paired-end reads on an SP flow cell for **Haplosclerida sp. nov**. and MiSEQ Micro 300-Cycle sequencing for *Xestospongia deweerdtae*. De novo genome assemblies were generated using SPAdes (version 3.15.5, Prjibelski *et al*., 2020) using the “—careful” option.

Contigs containing mitochondrial DNA, the *cox2* gene, and nuclear ribosomal RNA genes (*18S* and *28S*) were identified using the FASTA tool (Pearson and Lipman, 1988) by sequence similarity to homologous genes from the *Amphimedon queenslandica* genome (Erpenbeck *et al*., 2007; Srivastava *et al*., 2010). The primary mitochondrial contig was annotated using the MFannot web server (Lang *et al*., 2023). Additional transfer RNA (tRNA) genes were identified using tRNAscan-SE v2.0.6 (Lowe and Chan, 2016; Chan *et al*., 2021) in Organellar Mode (-O), enabling maximum sensitivity.

### Phylogenetic and correspondence analyses

*18S* and *28S* sequences of other haplosclerids were downloaded from GenBank. Alignments were made using MAFFT (version 7.508, Katoh and Standley 2013) with the “--auto” option, and highly variable sites were removed using GBlocks (Castresana 2000, -b5 = a, other settings default). The final alignment contained 880 positions for *18S* and 729 positions for *28S*. Maximum likelihood (ML) trees were constructed using RAxML-NG (version 1.1.0, Kozlov *et al*., 2019) using the GTR+G model, using 1000 bootstrap replicates.

Correspondence analysis was performed using the R package “ade4” version 1.7-22 (Dray and Dufour, 2007).

## Results

### Haplosclerida sp. nov. is morphologically distinct from *Haliclona plakophila*

**Haplosclerida sp. nov**. and *Haliclona plakophila* exhibit similar morphologies: growing as a thin veneer of tissue that coats the *Plakortis* basibiont with a patchy distribution that is easily detachable. In addition, the color, consistency, and lack of visible oscula make it almost impossible to distinguish the two epibiont species in situ (Figure 1). However, **Haplosclerida sp. nov**. can be recognized on the microscopic level. The ectosomal reticulation of **Haplosclerida sp. nov**. is mainly unispicular with a regular distribution of triangular meshes (Figure 2B). This contrasts with triangular meshes in *H. plakophila*, which vary between unispicular and paucispicular reticulation with some disorganization of the triangular meshes (Figure 2A). Oxea megascleres in **Haplosclerida sp. nov**. are larger and thicker (250 – 316 µm and 6 – 14 µm, respectively, from n=30 measurements) compared to those of *H. plakophila* (199–277 µm length; 3–9 µm thickness) (Figure 2C-D). In addition, the skeleton of **Haplosclerida sp. nov**. contains sigmas measuring 17–79 µm in length x 6–15 µm in width x 2–4 µm in thickness and exhibiting two different shapes (n=15) (Figure 2B).

**Figure 2.**
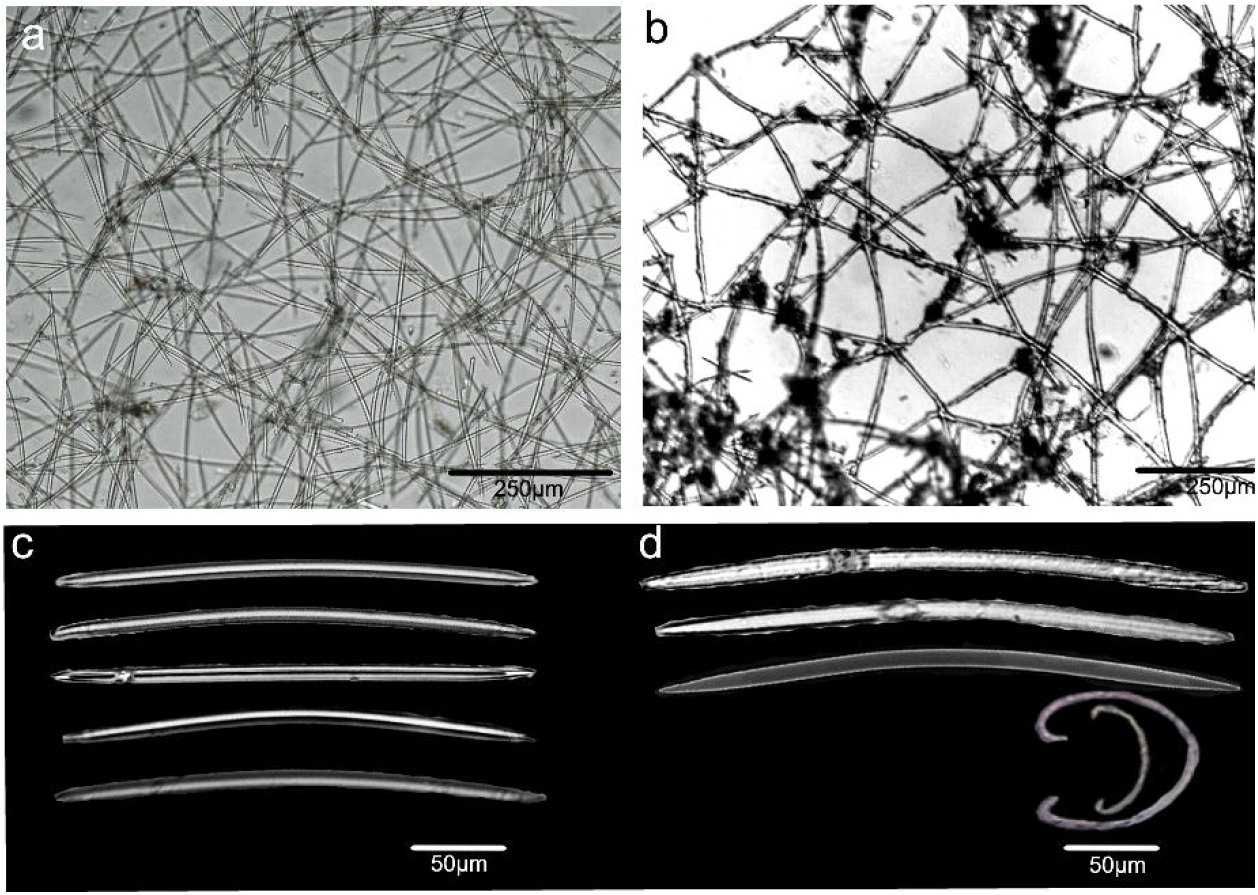
Ectosomal skeleton reticulation and spicule composition of *Haliclona (Halichoclona) plakophila* holotype USNM1254650 (A, C) and **Haplosclerida sp. nov**. BPBM C1914 (B, D). Spicule images are from light microscopy and scanning electron microscopy (C, D).

**Figure 3.**
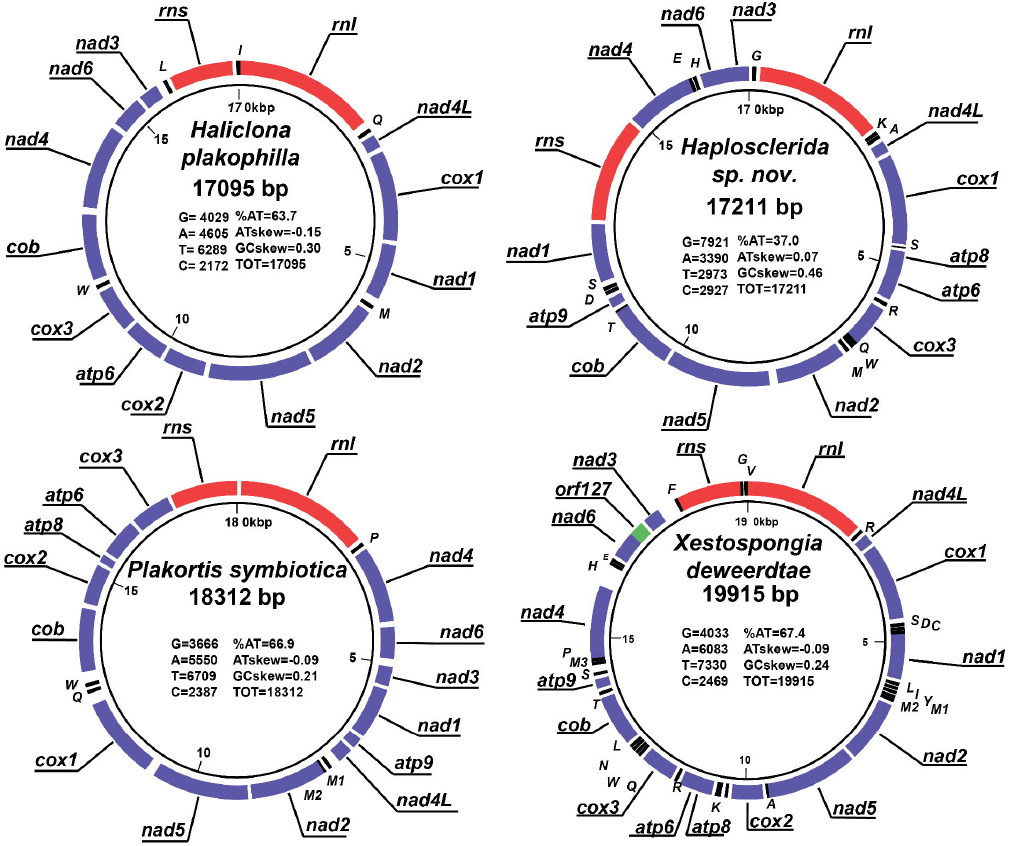
The mitochondrial genomes of the four sponges. Protein-coding genes are shown in purple, RNA genes in red, tRNAs in black, and intergenic regions in white. *atp* = ATP synthase subunits, *cox* = cytochrome oxidase subunits, *nad* = NADH dehydrogenase subunits, *cob* = apocytochrome b, *rnl* = large subunit rRNA, *rns* = small subunit rRNA. tRNAs are represented by their standard single-letter notation and anticodon in parentheses. The *orf127* in *X. deweerdtae* mitogenome is shown in green.

### Haplosclerida sp. nov. contains a highly unusual mitochondrial genome

The mtDNA of **Haplosclerida sp. nov**. is reconstructed as a 17.2-kbp-long circular molecule containing genes for 13 mitochondrial proteins (*atp6, atp8, atp9, cob, cox1, cox3*, and *nad1-nad6* plus *nad4L*), large and small ribosomal RNA subunits, but only 13 tRNAs (the minimum number of tRNA genes required for mitochondrial translation in demosponges is 24). *Atp8* and *atp6* overlap by one nucleotide, while *nad5* and *cob* are fused in frame, with *nad5* lacking a termination codon. The mtDNA nucleotide composition is highly biased, with 63% G+C content. The AT and GC skews are 0.07 and 0.46, respectively. *Cox2* is absent from the main contig but is found on a separate contig. The *cox2* gene has a %G+C content of 55%, and AT and GC skews of -0.04 and 0.38, respectively, comparable but less extreme than the main mitogenome. However, the noncoding regions of the remainder of the *cox2* contig have a %G+C content of 39.4%, and AT and CG skews of -0.004 and 0.042, respectively.

The high G+C content, combined with strong GC-skew, of the main mitogenome resulted in the coding sequences being highly enriched in Gs, especially at the third codon positions. Conversely, the proportion of Ts at the third position is unusually low (Figure 4). Similar, but less extreme biases were present at other codon positions, resulting in a highly unusual amino acid composition of encoded proteins. The same bias of G-richness at all codon positions and T-poorness at the 3^rd^ codon position was observed in the *cox2* gene (supplementary figure 1).

**Figure 4.**
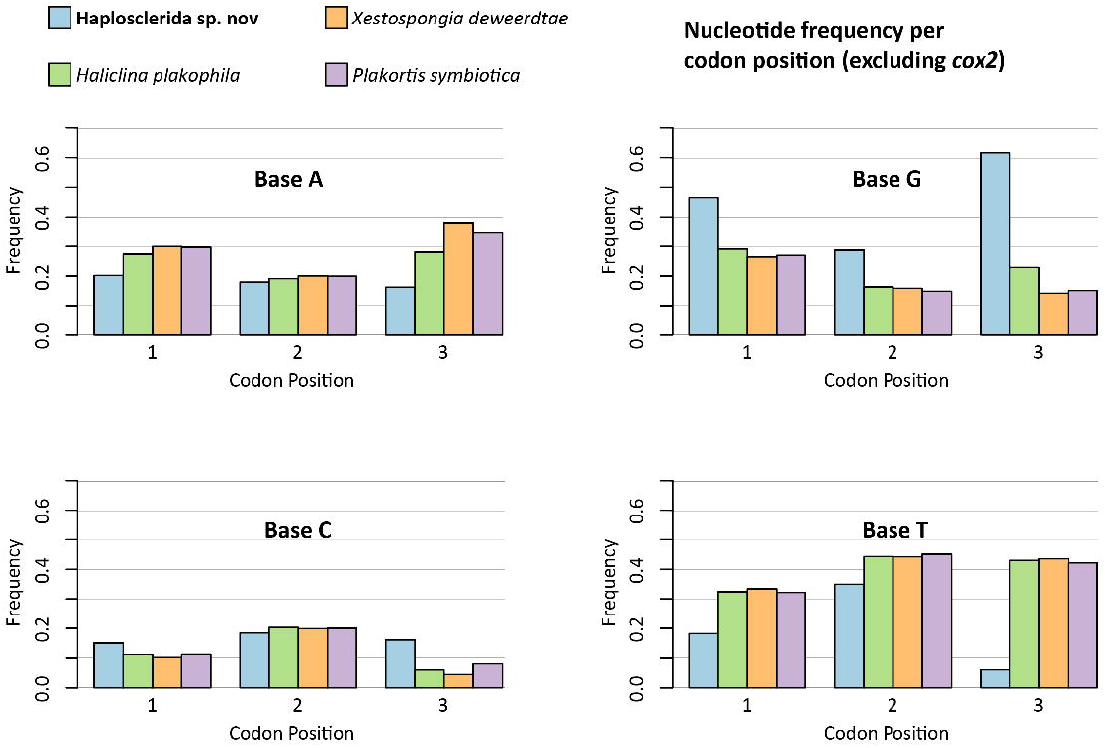
The frequency at which each nucleotide appears at each codon position in the coding sequences of the four sponges. There is a strong G-bias in the **Haplosclerid sp. nov**. mitogenome, especially at the third position, seemingly at the expense of Ts at the third position.

To check for possible misassembly as well as to infer the localization of *cox2*, we compared read coverage for genes in the assembled mitogenome, *cox2*, and short exonic portions of nuclear-encoded mitochondrial proteins, NADH dehydrogenase [ubiquinone] flavoprotein 1 (NDUFV1, %A+T ∼ 57%) and ATP synthase subunit alpha (ATPα, %A+T ∼ 53%). The average, per base, sequence coverage of *cox2* was 14.5, while the main mtDNA contig has an average coverage of 80. The average coverage for the first 200bp of NDUFV1 and the first 300bp of ATPα is ∼13.

The 13 tRNA genes range from 67 to 82 bp long (mean 71.2 bp) and show well-conserved secondary structures. The Infernal scores for the tRNAs were relatively low, ranging from 17 to 36.8, likely due to elevated GC content. The tRNA(Leu_CAA_) and tRNA(Leu_TAG_) are the longest at 82 bp, and both contain a well-conserved primary structure and a secondary structure that features a class I variable arm of 10 bases. The only other haplosclerids to have a reduced number of tRNA genes all belong to Clade B (Lavrov, Turner, and Vicente, 2025), suggesting that this is an independent origin of mt-tRNA gene loss among Haplosclerida.

### *Xestospongia deweerdtae* mitochondrial genome

A contig containing the *Xestospongia deweerdtae* mitogenome was recovered from the Spades assembly of low coverage DNA-seq data for this species and is reconstructed as a circular molecule, 19.9 kbp long, with 14 genes for mitochondrial proteins (*atp6, atp8, atp9, cob, cox1-cox3, nad1-nad6* plus *nad4L*), large and small subunit rRNA genes, and 23 tRNA genes. The genome has a %A+T content of 67.4%, and AT and GC skews of -0.09 and 0.24, respectively. The mitogenome also contains an open reading frame (orf127) that may code for a small ribosomal subunit protein, based on BLAST similarity results (E-value ≤ 0.001 compared to mitochondrial 28S ribosomal protein S24 found in animals such as *Drosophila*, E-value = 6e-04 compared to an uncharacterized protein in *Amphimedon queenslandica*). If this is confirmed, it would be the first reported ribosomal protein encoded in an animal mitogenome.

### Phylogenetic Analysis of nuclear rRNA data places **Haplosclerida sp. nov**. in a new clade within Haplosclerida

To clarify the phylogenetic position of **Haplosclerida sp. nov**., we extracted gene sequences for the *18S* and *28S* nuclear ribosomal DNA from DNAseq assembly. Phylogenies constructed using these sequences place the new species outside the five recognized clades of Haplosclerida (Redmond *et al*., 2013). However, the exact position of **Haplosclerida sp. nov**., as well as the interrelationship among the clades A, B, and D, is unresolved and differs between *18S* and *28S* analyses (Figure 5). Maximum Likelihood reconstruction using *18S* places this clade as a sister lineage to Clade D with weak support, while *28S* ML reconstruction places its sister to Clade B, also with weak support.

**Figure 5.**
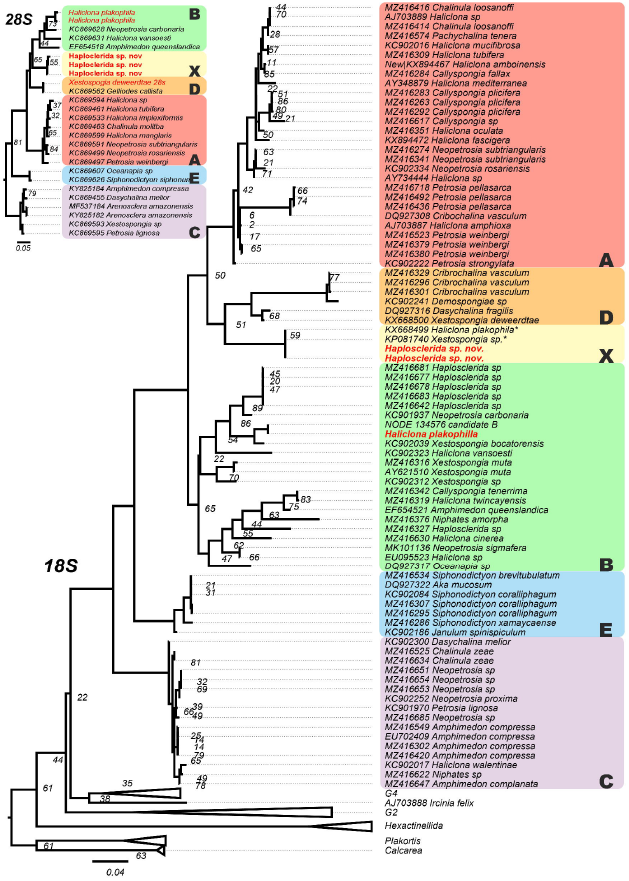
ML Phylogenetic trees using *18S* and *28S* genes. Haplosclerid clades are highlighted; the clade labeled X represents a new major clade not previously named. Sequences labeled with an asterisk denote previously reported sequences that were erroneously attributed to *Haliclona plakophila*, now recognized as belonging to **Haplosclerida sp. nov**. Sequences generated for this study are highlighted in red text. The *28S* tree is limited to only Haplosclerida. Bootstrap support values based on one thousand replicates are shown if less than 90%. No numbers indicate bootstrap support of > 90%.

### Mitochondrial codon and amino acid usage are highly biased in Haplosclerida sp. nov

We investigated how the highly unusual nucleotide composition of the **Haplosclerida sp. nov**. mitogenome influenced the amino acid compositions of the mitochondrial proteins. Similarly, given the location of *cox2* on a separate contig from the rest of the mitochondrial genes, we investigated patterns of codon usage between it and other mitochondrial genes. To achieve this, we performed two correspondence analyses (CA). The first CA assessed codon usage differences between the main mitochromosome and *cox2* (Figure 6A). The second CA evaluated differences in amino acid composition of mitochondrial proteins (excluding *cox2*) between **Haplosclerid sp. nov**. and other Haplosclerid sponges (Figure 6B).

**Figure 6.**
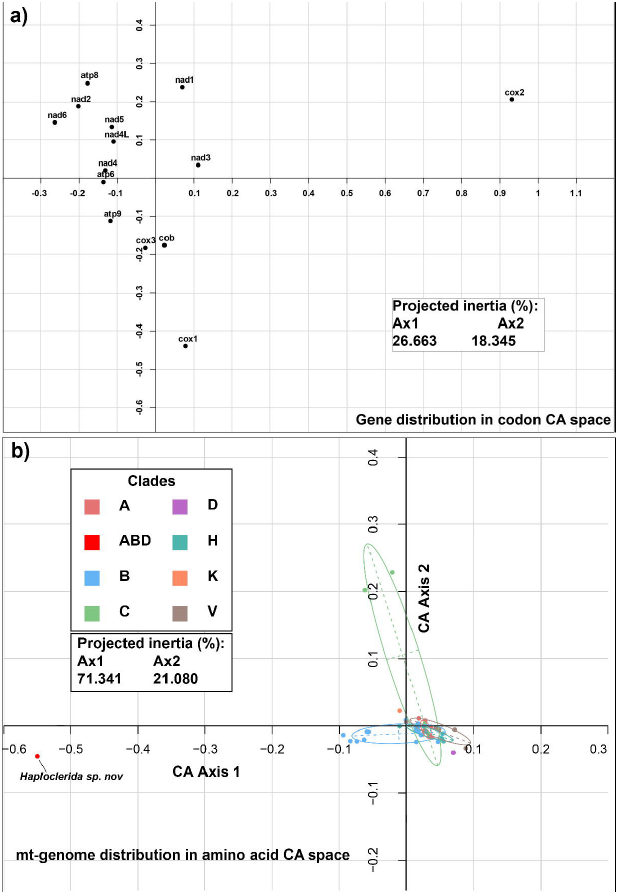
Correspondence analysis of (A) the codon usage between all genes within the mitogenome of **Haplosclerida sp. nov**. and the *cox2* gene, and (B) the amino acid composition of the mitochondrial genomes of haplosclerid sponges (excluding *cox2*).

The first CA shows that codon usage in protein-coding genes on the major mitochromosome of **Haplosclerida sp. nov**. cluster near one another, indicating a similar codon composition. However, *cox2* showed a large deviation in one axis. The X-axis explains 26.66% of the variation, and the Y-axis is 18.34%, together representing 45% of the variation. This suggests that the codon usage in *cox2 is* substantially different from the rest of the mitogenome.

The second CA shows that the amino acid usage of **Haplosclerida sp. nov**. is quite different from other haplosclerid sponges. Amino acid use is typically well clustered by clade, with a large variation being visible in the Y-axis in Clade C. However, **Haplosclerida sp. nov**. showed a large deviation away from the clusters on the X-axis. In this analysis, the X-axis explains 71.34% of the variation, and the Y-axis 21.08%, together accounting for 92.42%.

## Discussion

### A Third Symbiotic Partner of *Plakortis symbiotica*

**Haplosclerida sp. nov**. was first discovered through the detection of its unusual mitochondrial DNA within the symbiotic association between *Haliclona plakophila* and *Plakortis symbiotica*. Here, it is described as a novel species, distinguishable from the superficially similar *H. plakophila* by its spicule morphology and skeletal arrangement. So far, both **Haplosclerida sp. nov**. and *H. plakophila* have only been found as epizoic partners of *P. symbiotica*, with no evidence of a free-living lifestyle. By contrast, *Xestospongia deweerdtae*, the third haplosclerid observed to grow on *P. symbiotica*, is also found as an epibiont of *P. deweerdtaephila* and as a free-living sponge (Vicente *et al*., 2014). Our phylogenetic reconstructions show that the three epibionts are only distantly related, with **Haplosclerida sp. nov**. belonging to a new major clade of haplosclerids. These findings suggest a complex and diverse system of symbiotic relationships between *Plakortis* and haplosclerid sponges with at least three independent origins of specialized symbiotic associations involving *P. symbiotica*.

### Does unusual mitochondrial-nucleotide composition indicate unusual biology?

**Haplosclerida sp. nov**. has an unusual mitochondrial genome composition, with the coding strand being G-rich and T-poor. Consequently, this species also has a highly derived amino acid composition, manifested in a long branch on phylogenetic trees based on mitochondrial protein sequences and a large distance on the CA graph between this species and other haplosclerids. The unusual nucleotide composition is unlikely to be the result of positive selection on protein functions because the biases are strongest at the third codon position. Instead, we propose that the unusual nucleotide composition reflects the new species’ unusual biology, its likely obligatory symbiotic lifestyle, and possibly a parasitic relationship with its host sponge. Such a relationship could result in reduced locomotory demands on the larval stage, smaller effective population size, and host-associated bottlenecks, which, in turn, can lead to accumulation of deleterious mutations both in the mitochondrial and the nuclear genomes, including loss of genes involved in mitochondrial DNA repair. Indeed, a recent review linked parasitism in animals with higher mitochondrial evolutionary rates (Jakovlić *et al*., 2023).

The three main sources of mutations in animal mitochondria are replication errors made by DNA polymerase γ (*POLG*), spontaneous deamination of nucleotides in the leading strand displaced by the replication fork, and oxidative damage by free radicals created by electrons escaped from the electron transport chain (ETC) (Iliushchenko *et al*., 2025). Replication errors are predominantly C>T and A>G transitions. Because such errors are made during replication of both strands, they should not result in AT and GC-skews. Deamination of cytidine and adenosine also leads to C>T and A>G transitions, but is mostly limited to the leading strand during DNA replication and thus results in asymmetrical nucleotide composition between the two strands (Zheng *et al*., 2006). Unlike the other sources of mutations, oxidative damage to mtDNA most commonly produces G>T transversions caused by the oxidative adduct 8-oxo-guanine and contributes both to nucleotide skews and AT-richness of the mitogenome. Recently, it has been proposed that oxidative damage can also speed up adenine’s deamination, leading to the accumulation of Gs in the leading strand (Mikhailova *et al*., 2022; Iliushchenko *et al*., 2025).

While deamination of adenosine, either spontaneous or facilitated by oxidative damage, can lead to accumulation of guanines on the leading strand of mtDNA, it is expected to co-occur with deamination of cytidine, leading to accumulation of thymine. Indeed, in most of the mitogenomes, one of the DNA strands is GT-rich, while the other is AC-rich. Thus, the enrichment of Gs and the depletion of Ts in the coding sequences of **Haplosclerida sp. nov**. is unlikely to be explained by deamination. Also, a known defect in mtDNA repair, such as that in experimentally mutated *POLG* in mice, with reduced exonuclease activity (Trifunovic *et al*., 2004; Ameur *et al*., 2011; Maclaine *et al*., 2021) or the loss of mtDNA-encoded mutS homolog in octocorals (Muthye and Lavrov, 2021), would not explain strong strand asymmetry in the mtDNA of **Haplosclerida sp. nov**. Thus, the unusual nucleotide composition of **Haplosclerida sp. nov**. mitogenome remains a mystery.

### Is *cox2* a mitochondrial or a nuclear gene in **Haplosclerida sp. nov**.?

There is conflicting evidence regarding the localization of *cox2*, with some findings suggesting mitochondrial localization and others pointing to nuclear localization. Evidence supporting mitochondrial localization includes the absence of an identifiable N-terminal mitochondrial targeting sequence (MTS) in the encoded protein, as well as the G-rich and T-poor nucleotide composition of its coding sequence, similar to those of other mitochondrial genes. Furthermore, when translated using the standard genetic code, the *cox2* coding sequence contains an in-frame internal stop codon near the C-terminus, resulting in a polypeptide that is ten amino acids shorter than when translated using the minimally derived mitochondrial genetic code (code 4), previously inferred for demosponges.

In contrast, evidence favoring nuclear localization includes the assembly of *cox2* on a separate contig with lower sequence coverage and a codon usage pattern distinct from other mtDNA-encoded proteins, as revealed by correspondence analysis (CA).

However, none of these lines of evidence are definitive. On the one hand, multipartite mitogenomes have been observed in several animal groups, including ticks, nematodes, cnidarians, and calcareous sponges (Lavrov and Pett, 2016). More recently, multipartite mitochondrial genome organization has also been suggested for a species of demosponges (Paula *et al*., 2024). In some multipartite mitogenomes, different mitochondrial chromosomes show highly unequal sequencing coverage.

On the other hand, N-terminal MTSs are often absent in nuclear-encoded mitochondrial proteins, particularly in non-bilaterian animals (Muthye and Lavrov, 2018). Interestingly, the *cox2* coding sequence differs in nucleotide composition (e.g., %AT, AT skew, GC skew) from the surrounding sequences on the same contig. This contrast could suggest a recent transfer of *cox2* to the nucleus, with the sequence not yet equilibrated to match the nucleotide composition of its new genomic environment.

Interestingly, the translated COX2 polypeptide of **Haplosclerida sp. nov**. also shows slightly reduced hydrophobicity (∼47.9%) compared to other sponges in this study (∼51.5%), likely a result of its biased nucleotide composition. While not direct evidence of nuclear localization, this reduction in hydrophobicity could facilitate mitochondrial import of the protein if it is now nuclear encoded, a phenomenon previously observed in plants (Daley, Clifton and Whelan, 2002).

### Is this new species another *Haliclona*?

As a part of this study, we were interested in detecting morphological differences that would allow an easier distinction between **Haplosclerida sp. nov**. and *Haliclona plakophila*. Such differences were obtained both in the spicule composition and the ectosomal skeletal arrangement. A formal description with a comprehensive morphological examination will be provided in a future study. Here we discuss the overarching problem with naming this haplosclerid species.

The phylogenetic analyses conducted for this study place **Haplosclerida sp. nov**. in a new clade of Haplosclerida, suggesting the existence of a hitherto undescribed lineage of haplosclerids. The exact placement of this clade, however, is uncertain due to poorly resolved interrelationships among Clades A, B, and D (Figure 5). Since molecular estimates suggested that the most basal divergence within Haplosclerida occurred between 516 and 455 MYA (concurrent with the Cambrian Explosion, exceeding divergence times of most complex animal lineages), with the most basal divergence within the later-deriving Clade B occurring over 350 MYA (Lavrov *et al*., 2023; Lavrov, Turner and Vicente, 2025), it is clear that the species diverged from other haplosclerids hundreds of millions of years ago.

Despite observable morphological differences and a very distant phylogenetic placement from any other member of the genus, our initial assessment suggests that the new species fits the definition of the genus *Haliclona* Grant, 1841 as defined in the Systema Porifera as “Chalinidae with unispicular secondary lines” (De Weerdt 2002). *Haliclona* is a highly specious genus with 458 recognized species (de Voogd *et al*., 2025) and many more being described each year (Bispo, Willenz and Hajdu 2022, Bispo and Hajdu 2023, Vicente *et al*., 2025, Ott *et al*., 2024, van Soest 2024).

Molecular evidence from the past few decades has made it clear that the genus *Haliclona* is polyphyletic. The type species of the genus, *Haliclona (Haliclona) oculata* (Linnaeus, 1759), is reconstructed as a member of Clade A, while other *Haliclona*, for which molecular data are available, are placed in any of the Clades A, B, C and D (none have been recovered as members of Clade E, as far as we know). Nearly all other haplosclerid genera are also included in these clades.

The order Haplosclerida has an outstanding issue where morphologically derived taxonomy does not cooperate with molecularly reconstructed phylogeny. This creates a problem where, once officially described, the designation of species does not reflect their evolutionary origins and phylogenetic position. Indeed, most genera within Haplosclerida are polyphyletic, rendering their names as nothing more than convenient tools to facilitate a binomial nomenclature, while giving an impression of evolutionary relationships where none exist (McCormack, Erpenbeck and Van Soest 2002).

The unique phylogenetic position occupied by **Haplosclerida sp. nov**. and its ancient divergence time from other known species are grounds for the establishment of a new genus. However, classical taxonomic guidelines will result in this species, along with many other new species, being classified as *Haliclona*. There is an urgent need to reexamine the taxonomic guidelines by which Haplosclerida are classified. Here we continue to refer to this species as **Haplosclerida sp. nov**.

### Comparison with *Haliclona plakophila*: appearance and choice of host

**Haplosclerid sp. nov**. and *Haliclona plakophila* share a striking similarity in their external morphology and habit, both growing within the tissue of the basibiont *Plakortis symbiotica*, which protrudes through the surface, appearing as thin, bluish-white veneers of tissue. During our sampling expedition, it was noticed that the sample later identified as **Haplosclerida sp. nov**. appeared to protrude slightly above the surface of *Plakortis* basibiont compared to other samples. This veneer was also more readily peeled off. Interestingly, a similar “morphotype” was previously reported by Vicente, Zea, and Hill (2016) while describing the morphology of *Haliclona plakophila*. However, because we were only able to examine one specimen of **Haplosclerida sp. nov**., it is unclear if these features are typical of the new species and if the sample described by Vicente, Zea, and Hill contained a mixture of the two species. It is also possible that there are other distinctions in the morphology or lifestyle between the two species, which will be discovered by more careful examination.

The shared characteristics of **Haplosclerida sp. nov**. and *Haliclona plakophila* suggest new investigations in the ecology, geographic distribution, and life cycles of these sponges. It is generally believed that highly similar species rarely coexist without some form of differentiation in their ecology. They tend to evolve niche partitioning, trait divergence, or spatial/temporal strategies to avoid direct competition (den Boer, 1986). Where these differences do not exist, competitive exclusion or priority effects usually determine which species persist. It is unclear whether the predominance of *H. plakophila* among the sampled associations is an indication of its competitive advantage over **Haplosclerida sp. nov**. or the result of the priority effect for the sampling location. It should be informative to investigate these associations in other Caribbean locations.

It would also be interesting to investigate the biology of *Plakortis symbiotica* and the reason(s) this particular species is involved in several symbiotic relationships with haplosclerids. Species in the genus *Plakortis* (Plakinidae sponges) are usually well chemically protected against fouling by other organisms and are fierce allelopathic competitors. *P. symbiotica* does not appear to be an exception and is well defended against both fish and arthropod predators (Marty *et al*., 2017). Interestingly, *P. deweerdtaephila* and *P. symbiotica* have never been found in a free-living form despite considerable survey effort across the Caribbean (Vicente *et al*., 2014). Because Haplosclerida epibionts not only grow across the surface of their *Plakortis* basibiont but also within the mesohyl and can form deep oscular channels, it has been suggested that this may increase the pumping capacity of the basibiont (Vicente *et al*., 2014). However, this hypothesis has not been tested.

**Haplosclerida sp. nov**. represents a compelling addition to our understanding of sponge symbiosis and evolution. Its distinctive mitochondrial DNA and symbiotic lifestyle position it as a valuable model for exploring mitochondrial sequence evolution, host-symbiont interactions, and mechanisms of self- and non-self-recognition in sponges. Continued sampling and investigation will be essential to uncover the full extent of its biological significance and evolutionary history.

## Funding

This work was supported by the Presidential Interdisciplinary Research Seed (PIRS) Grant from Iowa State University to D.V.L. Postdoctoral work of J.V. was supported by the Division of Ocean Sciences at the National Science Foundation (OCE No. 2048457).

## Acknowledgement

Milton Carlo and Vance Vicente are thanked for their logistical support. The ToBo lab at HIMB and the Bernice Pauahi Bishop Museum are thanked for providing bench space for the morphological assessment of species.

## Notes

### Competing Interest Statement

The authors have declared no competing interest.

