## Supplementary figures and images for "Hidden in Plain Sight: A Novel Symbiotic Haplosclerid Sponge Species Revealed by its Mitochondrial Genome"

### SupFig1 - cox2CodonPosition.png

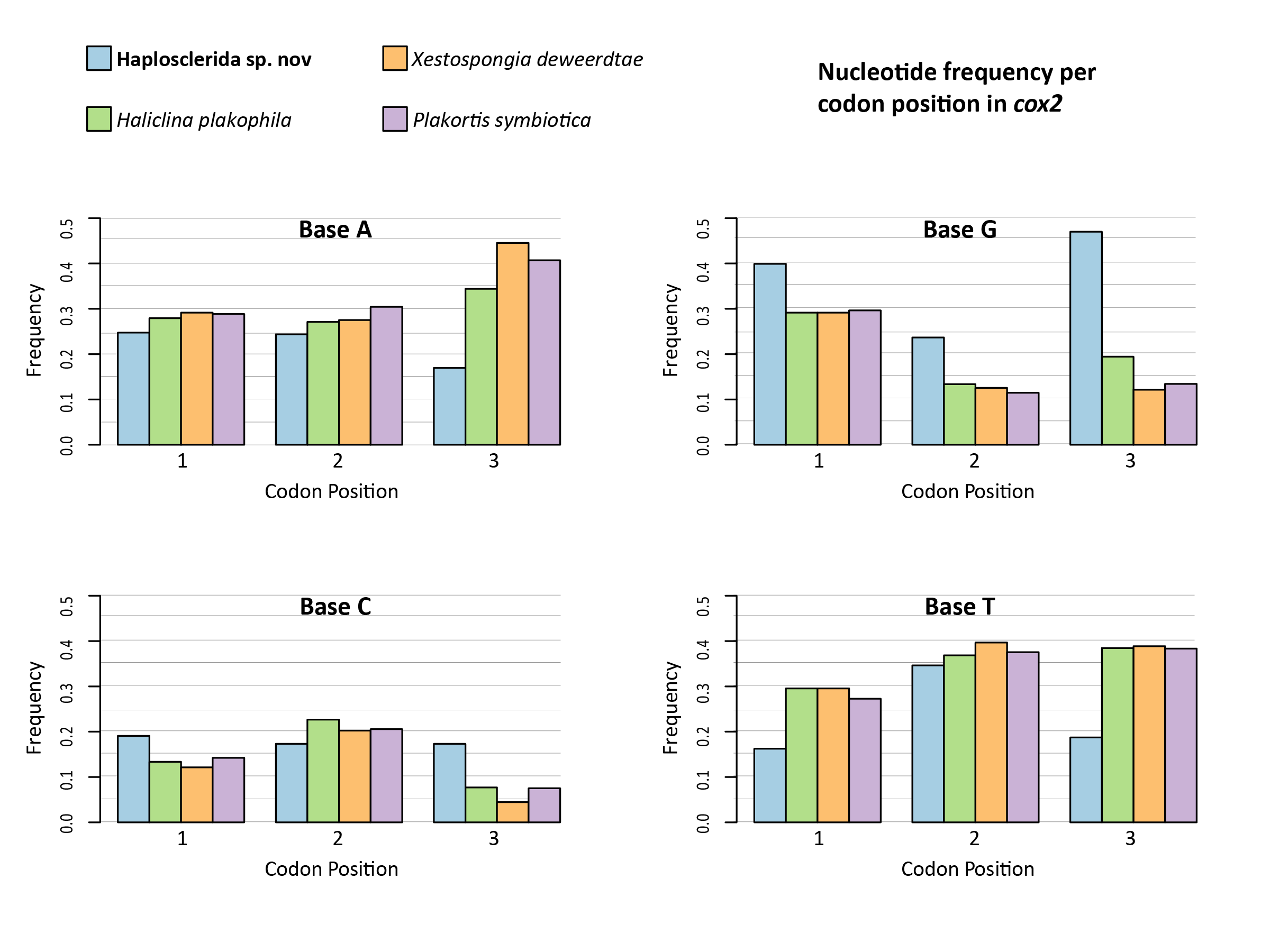
